# A core transcriptional signature of human microglia: derivation and utility in describing region-dependent alterations associated with Alzheimer’s disease

**DOI:** 10.1101/308908

**Authors:** Anirudh Patir, Barbara Shih, Barry W. McColl, Tom C. Freeman

## Abstract

Growing recognition of the pivotal role microglia play in neurodegenerative and neuroinflammatory disorders has accentuated the need to better characterize their function in health and disease. Studies in mouse, have applied transcriptome-wide profiling of microglia to reveal key features of microglial ontogeny, functional profile and phenotypic diversity. Whilst similar in many ways, human microglia exhibit clear differences to their mouse counterparts, underlining the need to develop a better understanding of the human microglial profile. On examining published microglia gene signatures, little consistency was observed between studies. Hence, we set out to define a conserved microglia signature of the human central nervous system (CNS), through a comprehensive meta-analysis of existing transcriptomic resources. Nine datasets derived from cells and tissue, isolated from different regions of the CNS across numerous donors, were subjected independently to an unbiased correlation network analysis. From each dataset, a list of coexpressing genes corresponding to microglia was identified. Comparison of individual microglia clusters showed 249 genes highly conserved between them. This core gene signature included all known markers and improves upon published microglial signatures. The utility of this signature was demonstrated by its use in detecting qualitative and quantitative region-specific alterations in aging and Alzheimer’s disease. These analyses highlighted the reactive response of microglia in vulnerable brain regions such as the entorhinal cortex and hippocampus, additionally implicating pathways associated with disease progression. We believe this resource and the analyses described here, will support further investigations in the contribution of human microglia towards the CNS in health and disease.

**Table of Contents:** Main points

- Published microglial transcriptional signatures in mouse and human show poor consensus.
- A core transcriptional signature of human microglia with 249 genes was derived and found conserved across brain regions, encompassing the CNS.
- The signature revealed region-dependent microglial alterations in Alzheimer’s, highlighting susceptible CNS regions and the involvement of TYROBP signaling.

## Introduction

Microglia are the most abundant myeloid cell type in the central nervous system (CNS), accounting for approximately 5-20% of the brain parenchyma depending on region (Lawson, Perry, Dri, & Gordon, 1990; Mittelbronn, Dietz, Schluesener, & Meyermann, 2001). These cells are phenotypically plastic and exhibit a wide spectrum of activity influenced by local and systemic factors (Cunningham, 2013; Perry & Holmes, 2014). Through development into adulthood, microglia influence the proliferation and differentiation of surrounding cells while regulating processes such as myelination, synaptic organization and synaptic signaling (Colonna & Butovsky, 2017; Hoshiko, Arnoux, Avignone, Yamamoto, & Audinat, 2012; Paolicelli et al., 2011; Prinz & Priller, 2014). As the primary immune sentinels of the CNS, microglia migrate towards lesions and sites of infection, where they attain an activated state that reflects their inflammatory environment (Leong & Ling, 1992). In these states, they can support tissue remodeling and phagocytose cellular debris, toxic protein aggregates and microbes (Colonna & Butovsky, 2017; Li & Barres, 2017). During neuroinflammation these cells coordinate an immune response by releasing cytokines, chemoattractants and presenting antigens, thereby communicating with other immune cells locally and recruited from the circulation (Hanisch & Kettenmann, 2007; Hickey & Kimura, 1988; Scholz & Woolf, 2007).

In common with mononuclear phagocyte populations throughout the body, recent studies have begun to reveal the diversity of microglial phenotypes in health, aging and disease states, as well as their unique molecular identity in relation to other CNS resident cells and non-parenchymal macrophages (Durafourt et al., 2012; Hanisch, 2013; Li & Barres, 2017; McCarthy; Salter & Stevens, 2017). The application of transcriptomic methods has been integral to these advances by enabling an unbiased and panoramic perspective of the functional profile of microglia. In addition to an improved understanding of the variety of context-dependent microglial phenotypes, other key benefits have arisen from these studies, notably the development of new tools to label, isolate and manipulate microglia (Bennett et al., 2016; Butovsky et al., 2014; Hickman et al., 2013; Satoh et al., 2016). Although most studies have been conducted in mice, a considerable body of data is now emerging from human post-mortem and biopsy tissue (Darmanis et al., 2015; Galatro et al., 2017; Gosselin et al., 2017; Olah et al., 2018; Y. Zhang et al., 2016). Whilst there are many conserved features between rodent and human microglia, the importance of further refining our understanding specifically of human microglia is underscored by important differences that have been observed between them (Butovsky et al., 2014; Galatro et al., 2017; Miller, Horvath, & Geschwind, 2010).

Recent transcriptomic studies have sought to characterize the human microglial transcriptomic signature from the CNS of non-neuropathologic individuals using data derived from either cells or tissue isolated from different brain regions (Darmanis et al., 2015; Galatro et al., 2017; Hawrylycz et al., 2012; Oldham et al., 2008). These analyses have been crucial in expanding our knowledge of their functional biology, however, our preliminary analyses found there to be little inter-study agreement across the published microglia gene signatures. Such inconsistency may have arisen due to technical differences in tissue sampling, brain areas analyzed, differences in patient characteristics and biological variance including the heterogeneity of different microglia populations (Grabert et al., 2016; Lai, Dhami, Dibal, & Todd, 2011; Lawson et al., 1990; Vincenti et al., 2016; Yokokura et al., 2011). This highlighted a need to derive a refined human microglial signature that would enable a more precise characterization of these cells in the healthy and diseased human brain. We therefore set out to define the core transcriptional signature of human microglia, i.e. shared by all microglial populations of the human CNS. To achieve this, we have performed an extensive meta-analysis of nine human cell and tissue transcriptomics datasets derived from numerous brain regions and donors. Secondly, we have used this signature to investigate region-dependent changes, while highlighting the influence of microglial numbers and activation in human tissue transcriptomics for Alzheimer’s and aging.

## Methods

### Comparison of published microglial signatures

Ten publications that defined microglial signatures, four in human and six in mouse, were identified (Table 1). To compare across studies, genes from each signature were converted to a common identifier i.e. HGNC (Povey et al., 2001) or MGI (Shaw, 2009) for human and mouse, respectively, using the online tool g:Profiler (Reimand et al., 2016). Subsequently, the tool was also used for interspecies comparison based on the MGI homology database, identifying human orthologues of mouse genes. At the time of analysis g:Profiler used Ensembl v89 and Ensembl Genome v36(Hubbard et al., 2002).

**Table 1.**
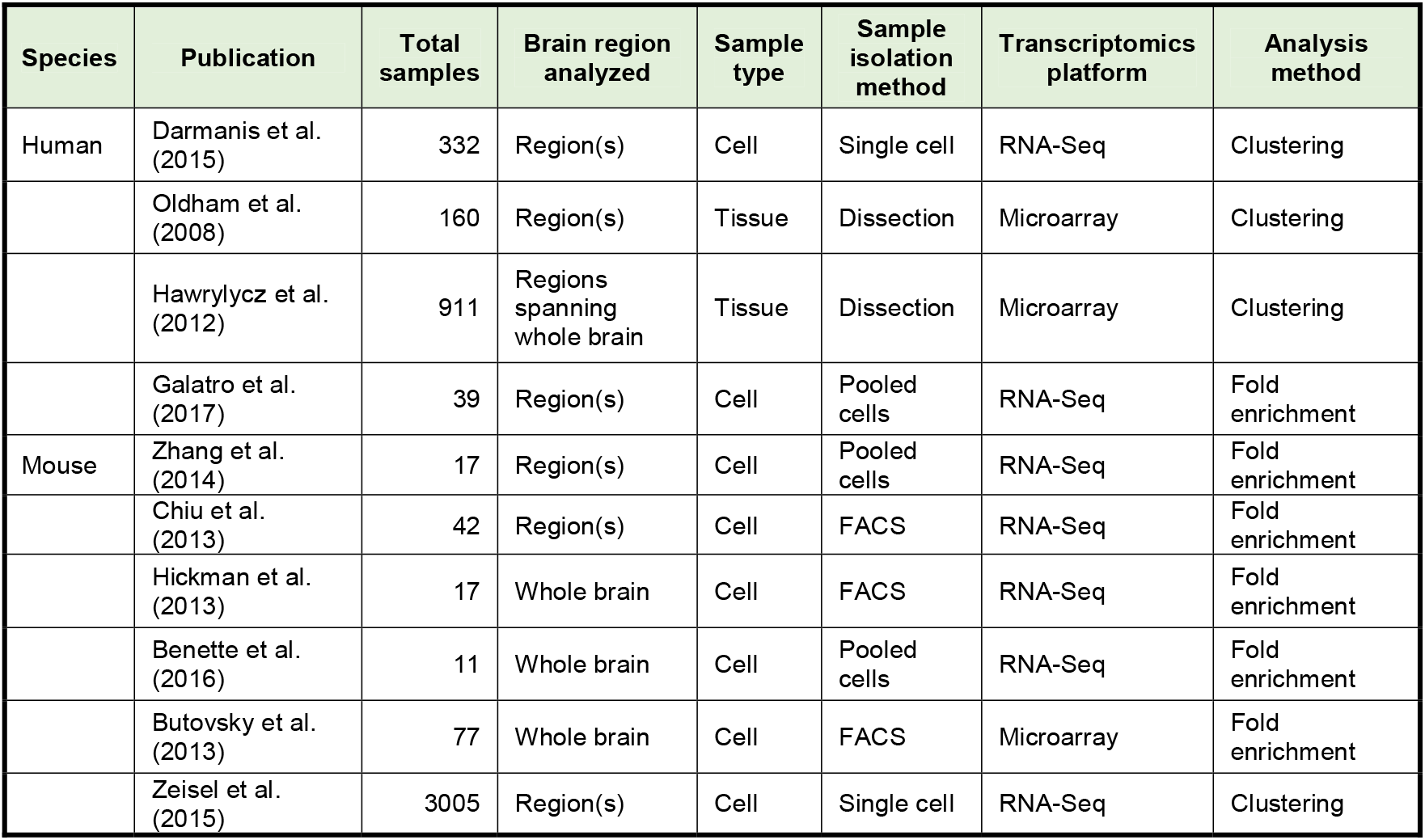
Experimental design of previous studies defining the microglial signature.

### Transcriptomics data acquisition and pre-processing

Tissue and cell transcriptomic datasets derived from the CNS were acquired for the derivation of the human microglial signature. These included data from the Genotype-Tissue Expression (GTEx) project (Lonsdale et al., 2013), Allen Brain Atlas (ABA)(Hawrylycz et al., 2012)(http://www.brain-map.org/) and from a study by Zhang *et al.*(Y. Zhang et al., 2016). The GTEx data comprised of two datasets, one generated on Affymetrix microarrays (n = 207) and a second by RNA-Seq (version 6, n = 1,259). In both cases, tissue samples were isolated from thirteen regions of the CNS at post-mortem, from individuals with no known neuropathology. The ABA data, generated on the Agilent microarray platform, consisted of 3,702 tissue samples taken from six individuals with up to 411 unique anatomical regions of the brain. Data from Zhang *et al.*(Y. Zhang et al., 2016) consisted of RNA-Seq data (n = 41) generated from different human CNS cell types (neuronal, glial and endothelial). For downstream analysis, data (n = 132) from immune (myeloid and lymphoid) and brain cell types (neuronal and glial) was downloaded (GSE49910) from the Gene Expression Omnibus (GEO)(Mabbott, Baillie, Brown, Freeman, & Hume, 2013). Lastly, for the analysis of microglia in aging and Alzheimer’s, data was derived from post-mortem samples of Alzheimer’s patients and controls from four cortical and hippocampal brain regions of 85 individuals (n = 235)(Berchtold et al., 2013). Further details of these datasets are provided in Table S1.

Transcriptomics data was downloaded from the appropriate sources and underwent stringent quality control. The unprocessed microarray data from GTEx (GSE45878) and the Alzheimer’s dataset (GSE48350), were downloaded from GEO. Data quality was assessed using the ArrayQualityMetrics package (Kauffmann, Gentleman, & Huber, 2008) in Bioconductor, and samples failing more than one of three metrics (between arrays comparison, array intensity distribution and variance mean difference) were removed. Subsequently, data was normalized using robust multiarray average from the oligo package (Carvalho & Irizarry, 2010) and where multiple probesets represented the same gene, the probeset with the highest average expression across donors was selected to represent the respective gene. GTEX and ABA preprocessed RNA-Seq data was downloaded directly from the GTEx portal and the ABA website, the latter consisting of six pre-normalized datasets from the brains of separate donors. Furthermore, quality control was conducted by inspecting sample-to-sample correlation networks using Graphia Professional (Kajeka Ltd, Edinburgh, UK), revealing outlier samples or batches effects. Evident from the GTEx RNA-Seq data, early batches (LCSET-1156 to LCSET-1480) poorly correlated with other samples, forming a highly connected group separate from other samples within the network. For downstream analysis, genes were filtered from the Affymetrix microarray, with a normalized expression level <20 and <1 FPKM or RPKM for RNA-Seq.

### Gene annotation through coexpression networks analysis

To define a core microglial gene signature, the tissue and cell transcriptomics datasets described above were analyzed using the coexpression network analysis tool Graphia Professional. For each dataset, Pearson correlations (*r*) were calculated between all genes to produce a gene-to-gene correlation matrix. From this matrix, a gene coexpression network (GCN) was generated, where nodes represented genes and genes correlating greater than a defined threshold were connected by edges. Coexpressed genes formed highly connected cliques within the overall topology of the graph, which were defined as clusters using the Markov clustering algorithm (MCL), with the default inflation value of 2.2 (Van Dongen, 2000). All Pearson threshold values used for individual datasets were above *r* ≥ 0.7 and thereby graphs included only correlations that were highly unlikely to occur by chance (Figure S1). For each dataset, the threshold for correlations was further adjusted to achieve a single microglial cluster containing the three canonical marker genes for microglia, *CX3CR1, AIF1* and *CSF1R* (Elmore et al., 2014; Mittelbronn et al., 2001). The final microglial gene signature was defined by genes present in at least three of the nine dataset derived microglial signatures (Table S2 and S3).

### Validation of the core human microglial signature

Various lines of evidence were investigated to validate the conserved nature of the derived human microglial signature. Firstly, the average expression of signature genes was compared between myeloid and other cell types from an atlas of primary human cells (Mabbott et al., 2013), using the Mann-Whitney U test. Similarly, the average expression of signature genes in the GTEx RNA-Seq data and donor one of the microarray ABA data was also compared with the microglial cell densities in mouse, for comparable regions (Lawson et al., 1990). Where available, immunohistochemical (IHC) staining of proteins encoded by signature genes were examined in the Human Protein Atlas (HPA)(Nilsson et al., 2005) across different regions of the human CNS. Enrichment analysis for Gene Ontologies (GO), pathways and transcription factor binding sites were conducted using ToppGene (Table S4)(Chen, Bardes, Aronow, & Jegga, 2009). In order to annotate the function of signature genes with relevance to myeloid and immune cells, the GeneCards database and literature were consulted (Table S3)(Safran et al., 2010).

Signature genes were then compared with other published mouse and human microglial signatures. The largest signature reported by Galatro *et al.* (Galatro et al., 2017), comprising of 1236 genes, included many genes from other microglia signatures and largely overlapped with the proposed signature. Therefore, to evaluate the specificity of the two signatures, we examined the expression of their respective genes in the GTEx RNA-Seq dataset. A GCN constructed (*r* ≥ 0.7) from the GTEx RNA-Seq dataset revealed five gene clusters enriched in Galatro *et al.* signature genes, representing various region-specific expression profiles (Table S5). Subsequently, for each cluster, the average expression of Galatro *et al.* genes was compared between the region with the highest expression versus the remaining samples, using the Mann-Whitney U test.

### Analysing microglia in aging and Alzheimer’s

To study microglia in aging and Alzheimer’s, samples from the study by Berchtold *et al.*(Berchtold et al., 2008) were binned into four age groups, 20-39, 40-59, 60-79 and 80-99 yr. The average expression level of microglial signature genes was calculated for samples in each age group and comparisons were made between Alzheimer’s samples (80-99 yr) with age-matched controls, and between older groups (80-99 yr) against younger control using the Mann-Whitney U test, corrected for multiple testing. To identify genes that represent microglial activation specifically in Alzheimer’s, a GCN (*r* ≥ 0.7) was constructed from only those samples derived from Alzheimer’s patients. The Fischer’s exact test was used to identify clusters enriched (adjusted *P* < 0.01) in core signature genes. Two such clusters were identified, clusters 5 and 67, containing 333 and 18 genes, respectively. Of these genes, 165 were not part of the derived signature and were considered as potential microglial associated genes (MAGs) in Alzheimer’s (Table S6). To aid the interpretation of the MAGs, their enrichment of pathways, GO annotations and associated transcription factor binding sites, were calculated using ToppGene (Table S7)(Chen et al., 2009). Following this, differentially expressed core genes and MAGs across brain regions, were identified for the superior frontal gyrus, between old (< 60 yr) and young controls (≥ 60 yr), and also between Alzheimer’s (≥ 60 yr) and age-matched controls, using the limma package in R (Table S6)(Smyth, 2005). A similar enrichment analysis was conducted for genes differentially expressed between Alzheimer’s samples and age-matched controls (Table S8).

## Results

### Heterogeneity of existing microglial signatures from human and mouse

To examine the human microglia gene signature, previous signatures from human brain tissue or cells were compared (Table 1, Figure 1). Four such studies varied considerably in the number of genes they defined, ranging from 21 to 1,236 genes. Of the 1,464 unique genes identified in all these studies, only a fraction (15%, 214 genes) were present in two or more signatures, with only 10 genes reported by all four publications. To verify that these results were not purely attributed by the individual variation in humans, the six publications reporting mouse-microglial signatures were also compared. Altogether these listed 690 genes (ranging from 47 to 433 genes) with 300 orthologues common to studies in human. Similar to the comparison of human signatures, only 26% (179 genes) of genes were reported by more than one study, with only 9 genes common to all. These observations highlight the discordance between existing microglia marker lists and a need to develop a robust and validated human microglial gene signature.

**Figure 1.**
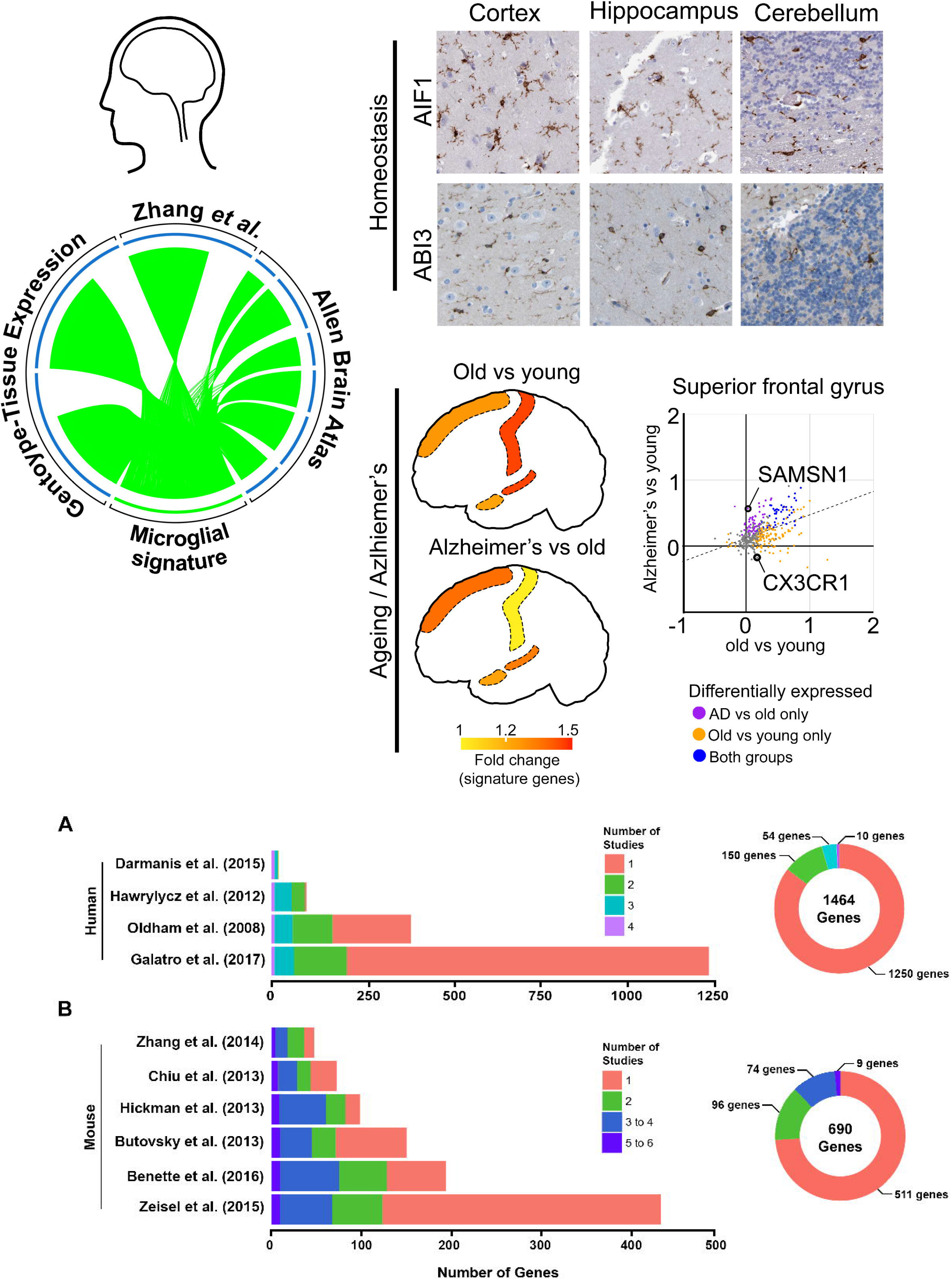
Comparison of published human and mouse microglial signatures. Comparison of signature size and gene overlaps amongst microglial signatures defined by studies in **(A)** human and **(B)** mouse (left panel). Plot (right panel) of gene overlap considering all microglial genes identified in all studies for the respective species

### Derivation of a conserved core human microglial signature

Observing the variability across published studies, we set out to define a human microglia gene signature from human tissue and cell data using a GCN (Figure 2A, Table S1). For this meta-analysis, tissue datasets including the GTEx project and ABA data were chosen, which cover a broad spectrum of sampling, across numerous CNS regions and donors (Hawrylycz et al., 2012; Lonsdale et al., 2013; Shen, Overly, & Jones, 2012). Additionally, Cell transcriptomic data from Zhang *et al.* reporting the top 20 marker genes for different brain cell types, was also included in the analysis (Y. Zhang et al., 2016). Post QC, the data amounted to a total of 5,020 samples isolated from 197 donors and 440 anatomical regions of the brain. To extract a microglial cluster from individual datasets, each was analyzed independently using a GCN (Tom C Freeman et al., 2007; Theocharidis, Van Dongen, Enright, & Freeman, 2009). This method exploits the inherent variability amongst samples due to variation in sampling, donors and cellular diversity across different CNS regions. In this case, genes expressed specifically by microglia in the context of the CNS, will vary in expression according to the regional abundance of these cells and therefore correlate in their expression, e.g. the poorly populated cerebellum presents a low expression of these genes relative to other regions. For constructing GCNs, genes are represented by nodes, and connected by an edge based on the similarity between their expression profiles, as quantified by the Pearson correlation coefficient (Figure S1). In this network, correlated genes form highly connected cliques within the overall topology of the GCN, and are defined as clusters using the MCL algorithm (Enright, Van Dongen, & Ouzounis, 2002; B. B. Shih et al., 2017). Using this approach on individual datasets, a microglial cluster containing the known marker genes *CX3CR1, AIF1* and *CSF1R*, was identified for each dataset (Table S2)(Elmore et al., 2014; Mittelbronn et al., 2001). The final high confidence microglia gene signature was defined by 249 genes, which were present in three or more dataset-derived clusters, so as to avoid biases towards individual datasets. However, it should be noted that the 395 genes observed in at least two dataset-derived microglial clusters also showed a strong enrichment for genes with a known immunobiological function (Table S3).

**Figure 2.**
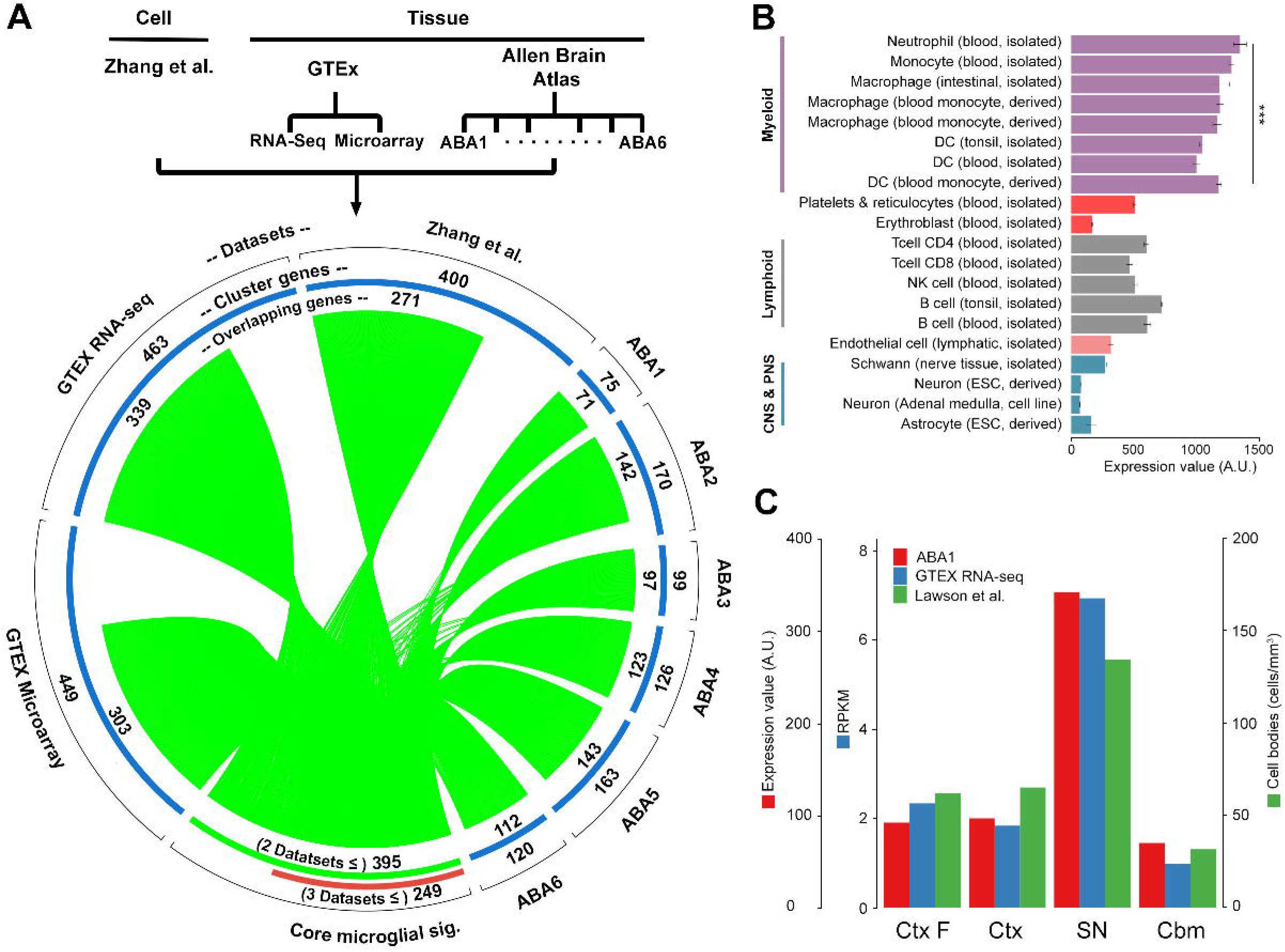
Microglial signature derivation, comparison and validation. **(A)** A diagram showing steps in the derivation of the core microglial signature. From each dataset (upper panel) a microglial-specific cluster was identified using coexpression network analysis (blue sectors). In comparing these gene clusters, 395 overlapped across more than one dataset (green sector). From this set of overlapping genes, green lines connect a specific gene to all datasets in which it was identified. Of the overlapping genes, those co-occurring in three or more datasets were taken to represent the core microglial signature (red sector). **(B)** Average expression of core signature genes in various neuronal and immune cell types selected from an expression atlas of primary cells (Mabbott et al., 2013). **(C)** Comparison of the average expression across tissue transcriptomics data and microglial cell numbers in mouse, for regions common to the respective studies (Lawson et al., 1990). Abbreviations - AU: Arbitrary units; sig: Signature; ABA: Allen Brain Atlas; Ctx F: Frontal cortex; Ctx: Cortex; SN: Substantia Nigra; Cbm: Cerebellum. *** significant at *P* < 0.001.

### Validation and description of the core human microglial signature

To validate the microglial signature genes, various lines of evidence were examined. First, a comparison of the average expression of core signature genes across cell types revealed a significantly higher (*P* < 0.001) expression in myeloid cells relative to other immune (most of which are scarce within non-neuropathologic brain tissue) and CNS cell types (Figure 2B)(Ginhoux et al., 2010). Second, the average expression of core genes across brain regions in the GTEx and ABA datasets correlated with regional microglial densities as measured in the mouse (Figure 2C)(Lawson et al., 1990). Third, where data was available, the IHC staining of proteins encoded by signature genes was examined in the CNS. This confirmed the microglial expression of known markers e.g. *AIF1*, as well as less characterized proteins in the core set, e.g. *APBB1IP, ABI3, FCER1G* and *ARHGDIB*, which specifically stained for microglia across the four regions analyzed by the HPA resource (Figure 3A)(Nilsson et al., 2005). Finally, GO enrichment analysis was performed and complemented by manual annotation of the core human microglial gene signature. Literature mining showed most genes in the list to have some association with microglial/macrophage biology and overall there was a significant enrichment in genes known to be associated with microglial processes (Table S3, 4). These include TLR signaling (*TLR1, TLR2*), complement pathway (*C3AR1, C1QA* and *C2*), TYROBP signaling (*TREM2, TYROBP*), and cytoskeletal organization (*AIF1, CAPG* and *WAS*) (Figure 3C)(Hong et al., 2016; Marinelli et al., 2015; Yeh, Hansen, & Sheng, 2017). Genes recently identified as highly enriched in human or mouse microglia, relative to other macrophages and CNS cells, were also present in the signature (e.g. *GPR34, P2RY12, P2RY13, TMEM119*)(Butovsky et al., 2014).

**Figure 3.**
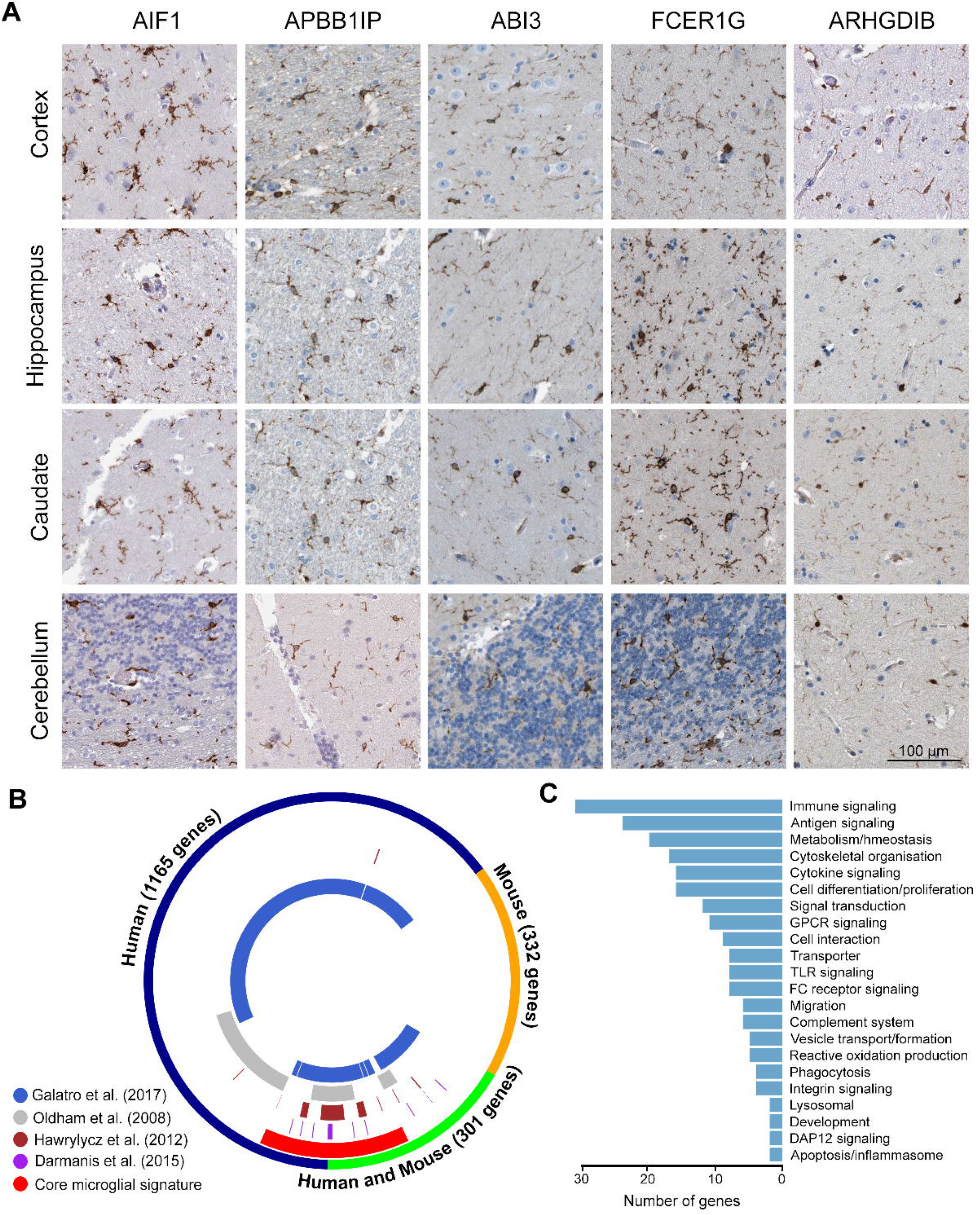
Supporting evidence for core microglial signature. **(A)** IHC staining of proteins of signature genes taken from the HPA, specifically staining for microglia within CNS sections from various regions (Nilsson et al., 2005). **(B)** Comparison of published human microglial signatures (inner circles) with reference to all the genes identified in both mouse and human studies (outer most circle), including the current core microglial signature (red segment). **(C)** Annotation of the derived microglial signature genes based on GeneCards, with relevance to myeloid and immune cells (Safran et al., 2010).

The core signature was then compared to the published microglial signatures from both human and mouse (Figure 3B). The majority of genes (248 genes) overlapped with signatures from earlier works, with *HLA-DRB3* being unique to this study. Over half of the core signature genes included those overlapping between published human and mouse signatures, while the remaining genes were specific to previous signatures in human (113 and 134 respectively). A majority of the core signature genes (64%, 142 genes) were identified in two or more human studies, whilst 99 genes overlapped solely with the Galatro *et al.* signature. To further validate the specificity of the current microglial signature, the coexpression of these genes was compared with that of the Galatro *et al.* signature (1,236 genes), which included the majority of genes in other signatures. On constructing a GCN from the GTEx RNA-Seq dataset, genes of the current signature were strongly coexpressed with one another within the network graph (Figure 4A). Many Galatro *et al.* signature genes were similarly coexpressed, however, many others were scattered across the network, indicative of an overall poor correlation between them in comparison to the current signature (Figure 4B). Cluster analysis showed a contrast in the expression profiles of clusters enriched in Galatro *et al.* signature genes relative to the microglial cluster as defined by marker genes (cluster 6, Figure 4C, D, and E). The expression pattern of these clusters deviated from the microglial cluster 6 and presented significant (FDR < 0.001) region-specific expression (Table S5). For instance cluster 1 containing 94 genes from the Galatro *et al.* signature were highly expressed in the cerebellum, a brain region having a low number of microglia. On comparing these genes with a recently published list of cerebellum-specific mouse microglial genes (Grabert et al., 2016), only three genes coincided and analysis of HPA IHC data suggested that whilst some were specifically expressed in microglia in other regions, they were not microglial specific in the cerebellum (Figure S2).

**Figure 4.**
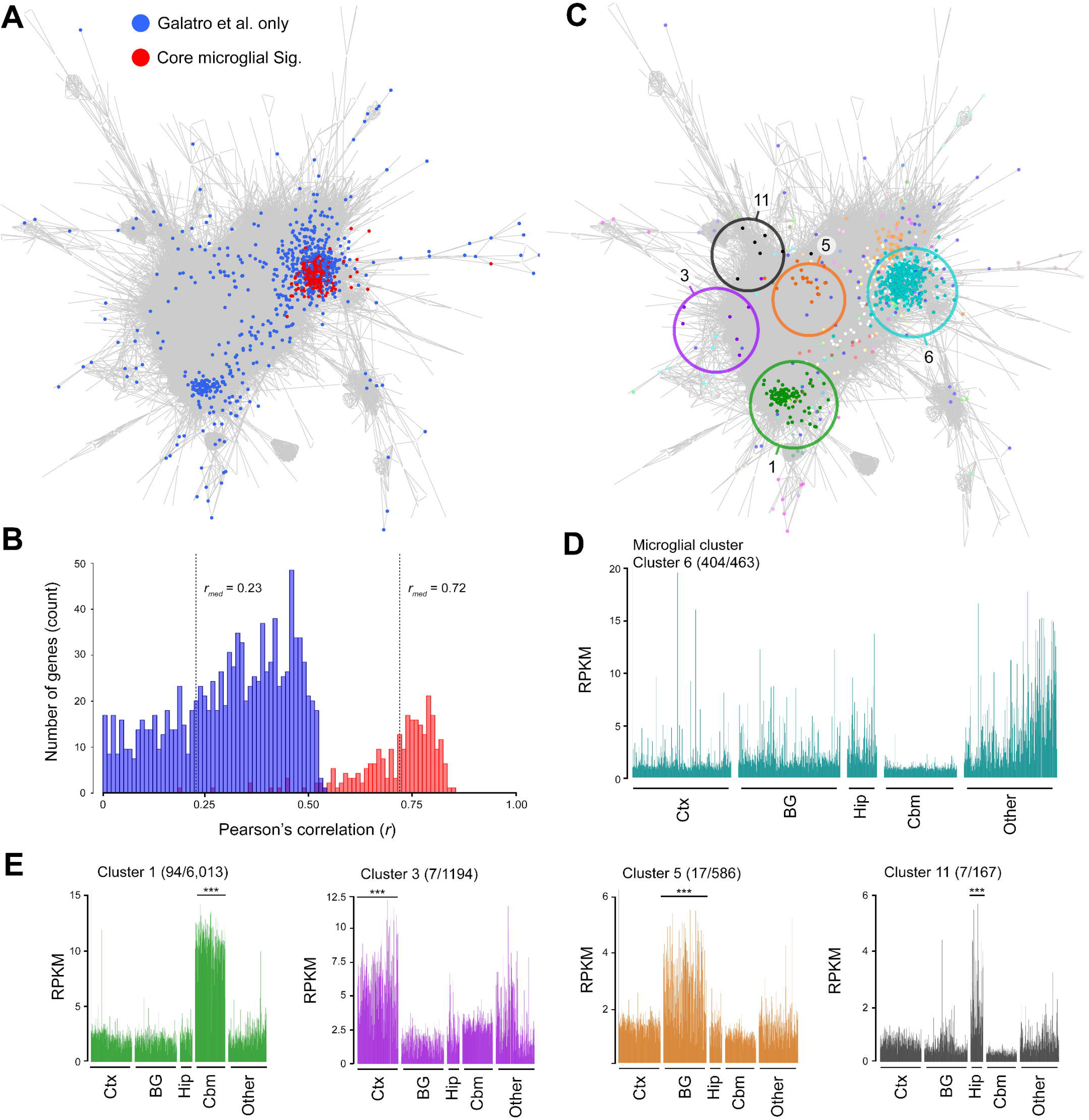
Coexpression of Galatro *et al.* signature and core microglial signature in the CNS. **(A)** Coexpression of Galatro *et al.* microglial signature (blue) and the current microglial signature (red) within the GTEx RNA-Seq data. **(B)** Histogram of the genes based on their median correlation with other genes of the same signature. Note the overall lower correlation between genes of the Galatro signature compared to the current signature. **(C)** Cluster analysis of the GTEx data, showing five clusters enriched in Galatro *et al.* signature genes. **(D)** Profile of cluster having 463 genes of which 404 are present in Galatro *et al.* signature, also containing majority of the current list. **(E)** Expression profile of Galatro *et al.* signature from the clusters identified. Abbreviations - Sig: Signature; Ctx: Cortex; BG: Basal ganglia; Hip: Hippocampus; Cbm: Cerebellum. *** significant at *P* < 0.001.

### Microglia in Alzheimer’s disease

We next used the 249 gene signature to assess the human microglial profile in aging and Alzheimer’s through analysis of a transcriptomics dataset derived from cortical and hippocampal regions of Alzheimer’s patients and non-neuropathic controls (Berchtold et al., 2013)(Table S1). As a preliminary analysis, the average expression of signature genes was used as a proxy measure of microglial number and calculated for all 20 yr age groups across regions (Figure 5A). Apart from the entorhinal cortex, a significant increase in expression of core genes was observed with aging. For example, in the hippocampus, a 1.6 fold change (FC) in expression (*FDR* < 0.01) was observed between the oldest and youngest control age groups. The lack of significance for the entorhinal cortex is likely attributed to the significant variation between samples across the different age groups. On comparing the average expression of core genes in Alzheimer’s with age-matched controls, the superior frontal gyrus showed a significant increase in Alzheimer’s samples (FC = 1.2, *FDR* < 0.05), a region known to be significantly affected in both aging and Alzheimer’s, based on neuronal connectivity studies (Bakkour, Morris, Wolk, & Dickerson, 2013; Stam, 2014). Although non-significant, other regions also showed a consistent increase in expression of the signature genes relative to age-matched controls.

**Figure 5.**
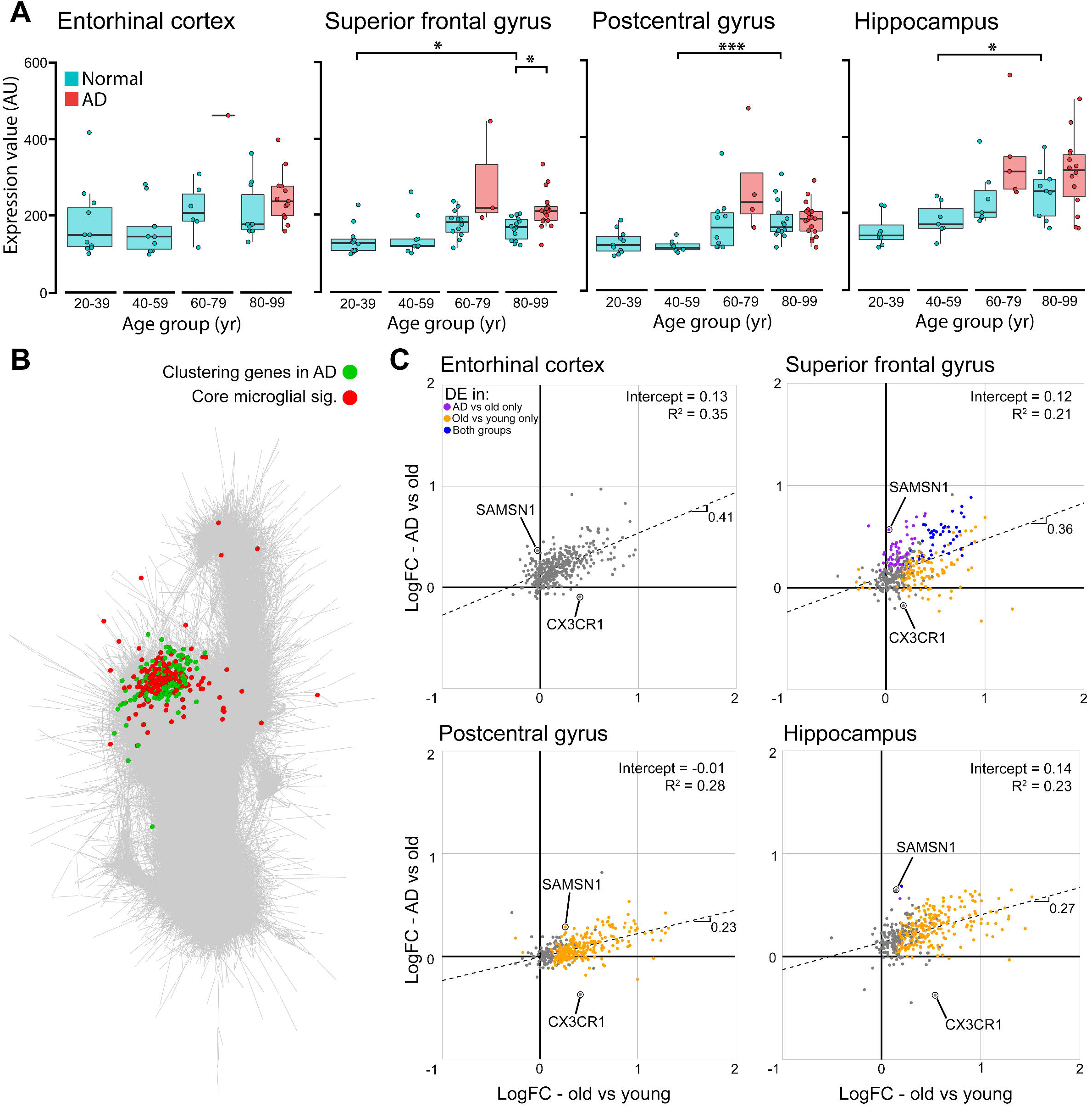
The microglial response to Alzheimer’s disease. **(A)** Average expression of core signature genes in normal and Alzheimer’s samples from different age groups. **(B)** Coexpression network highlighting core (red) and microglial associated genes (green) in Alzheimer’s samples. **(C)** Comparison between all the MAGs and core genes of the fold change in Alzheimer’s versus age-matched controls (y-axis) and between old versus young controls (x-axis), across regions. For each comparison differentially expressed genes are shown, in the process of aging (yellow), in Alzheimer’s (purple) or differentially expressed in both processes (blue). The trend in expression of these in Alzheimer’s and aging is represented by the regression line (dashed line) with their slope, intercept and R^2^. Abbreviations - Sig: signature; AD: Alzheimer’s disease. * significant at *FDR* < 0.05, *** significant at *FDR* < 0.001.

Based on the hypothesis that microglia in Alzheimer’s not only increase in number but are also phenotypically altered by the presence of misfolded beta-amyloid protein and other potential biochemical stressors, we sought to identify other genes which were specifically coexpressed with the core signature genes across in brain samples from Alzheimer’s patients (Manocha et al., 2016). A GCN was generated using only those samples derived from Alzheimer’s patients (*r* ≥ 0.7), and two clusters were found enriched in core microglial genes based on a Fisher’s exact test (adj. *P* < 0.01). The 165 non-core genes were also present in these clusters, i.e. coexpressed with the core genes and used for downstream analyses (Figure 5B, Table S6). Enrichment analyses of these MAGs conducted using ToppGene (Chen et al., 2009) revealed GO terms associated with cell activation, wound healing, angiogenesis, apoptosis and immune defense response (Table S7). These analyses were complemented by an enrichment in the MAGs for pathways linked to platelet activation, NFKB signaling, TGFB-SMAD signaling and VLDL metabolism. Additionally, an enrichment of the *ETS2* binding site was observed in these genes, a transcription factor implicated in Alzheimer’s and a known transactivator of the APP promoter (Wolvetang et al., 2003).

In order to identify quantitatively, genes specifically associated with microglia in Alzheimer’s but not aging, a differential expression analysis was conducted based on the MAGs and core genes, to compare the response of microglia in aging and Alzheimer’s. Thus, the expression fold change between the old (≥60 yr) and young, was compared with that of Alzheimer’s and age-matched controls (Figure 5C, Table S6). Reinforcing our preliminary analysis in estimating microglial numbers in Alzheimer’s versus age-matched controls, the majority of differentially expressed genes (FDR < 0.05) were restricted to the superior frontal gyrus. Interestingly, the trends in expression for each region (represented by the regression line) matched the degree to which each region undergoes neurodegeneration in Alzheimer’s, e.g. the post-central gyrus, which is comparatively unaffected in Alzheimer’s relative to other regions of the brain (Thompson et al., 2003), showed the least upward trend (intercept = −0.01, slope = 0.23). In contrast gene expression in the entorhinal cortex and hippocampus, regions considered vulnerable to Alzheimer’s showed an upward trend, highlighting the significance of these genes in Alzheimer’s and not only aging. Although the genes differentially expressed across regions were not all the same, certain genes such as *SAMSN1* (superior frontal gyrus: FC = 1.48, *FDR* < 0.003) and *CX3CR1* (superior frontal gyrus: FC = 0.88, *FDR* < 0.707) had a consistent expression pattern across regions when comparing the expression fold change in Alzheimer’s and aging. To better characterize the microglial response in Alzheimer’s, we focused on the superior frontal gyrus, having the most number of differentially expressed genes and a significantly affected region in Alzheimer’s. In identifying genes likely representing changes in activation state rather than cell number, we considered the 52 genes differentially expressed only in Alzheimer’s versus age-matched controls. Here, genes also differentially expressed in aged versus young were excluded as they are known to be influenced by microglia abundance. Enrichment analysis of these genes highlighted processes related to cell activation (*PYCARD* and *PIK3CG*), wound healing (*A2M* and *SERPING1*), innate immune response (*TLR5* and *ITGAM*), and pathways associated with phagocytosis, TLR cascade, and cell activation linked with neuronal survival (Table S8). Moreover, several members of TYROBP signaling pathway were differentially expressed (*SAMSN1, SIRPβ2, CD37, IL10RA, PIP3CG* and *BIN2*), a pathway dysregulated in microglia during Alzheimer’s (Keren-Shaul et al.; Ma, Jiang, Tan, & Yu, 2015; B. Zhang et al., 2013). Of the differentially expressed genes, eleven were MAGs including *LYZ, RPS6KA1* and *SLA*, with known associations to Alzheimer’s (Ellison, Bradley-Whitman, & Lovell, 2017; Hu, Xin, Hu, Zhang, & Wang, 2017; Tuppo & Arias, 2005). Interestingly, certain classical microglial marker genes were differentially upregulated in Alzheimer’s e.g. *ITGAM* and *PTPRC*, while others showed a downward trend, including *CX3CR1* and *P2RY12;* the latter consistent with a loss of homeostatic microglial genes observed in Alzheimer’s mouse models (Keren-Shaul et al.). Alternatively, whilst tissue gene expression can be influenced by cell activation and cell numbers, certain genes found differentially upregulated in both Alzheimer’s and aging, such as *TSPO, MS4A6A* and MHC class 2 genes, are known contributors of microglial activation based on previous studies (Bergen, Kaing, Jacoline, Gorgels, & Janssen, 2015; Hamelin et al., 2016; Hu et al., 2017). Overall, these observations demonstrate the value of the refined microglial signature we have derived in deducing changes in microglial profile (numbers and functional status) in Alzheimer’s and are consistent with the region-specific vulnerability and progression of Alzheimer’s pathology.

## Discussion

Recent transcriptomic studies, majority of which have been conducted in mice, have greatly advanced our knowledge of the functional profile of microglia (Butovsky et al., 2014; Darmanis et al., 2015; Galatro et al., 2017; Hickman et al., 2013; Zeisel et al., 2015), their regional heterogeneity in the CNS (Grabert et al., 2016), and altered profile associated with neurodegeneration (Keren-Shaul et al.; Miller et al., 2010; Vincenti et al., 2016). Additionally, key differences between mouse and human microglia have been suggested (Galatro et al., 2017; Olah et al., 2018), emphasizing the importance of better characterizing the functional profile of human microglia in health and disease. Our initial investigations demonstrated that published microglia gene signatures vary considerably in their size and composition relative to one another. Contributors of the observed discrepancy are likely the differing experimental objectives, p-value thresholds or fold-change enrichment in defining signature genes, donor variability, differing analysis platforms/methods, regions examined and cell isolation methodologies (Okaty, Sugino, & Nelson, 2011). Indeed, there appears to be little consensus amongst current studies over the functional profile of microglia beyond a few well-known markers, e.g. *AIF1, CSF1R* and *CX3CR1* (Elmore et al., 2014; Mittelbronn et al., 2001).

To identify a conserved human microglial signature, we used an unbiased correlation network analysis, harnessing the power of cell and tissue transcriptomics data including two large studies; namely the GTEx and ABA datasets. Together they provide the largest publicly available transcriptomic datasets covering a comprehensive range of brain regions, collected from numerous donors (Hawrylycz et al., 2012; Lonsdale et al., 2013). GCNs were constructed to identify groups of genes with similar expression profiles, corresponding to cells or pathways, as has been shown possible using this approach (Tom C. Freeman et al., 2012; Mabbott et al., 2013; B. B. Shih et al., 2017). From each dataset, a microglia cluster was identified, based on the presence of canonical marker genes for these cells. The consensus from these dataset derived signatures, provided 249 genes representative of human microglia across datasets. To our knowledge, this is the first study to deconvolute a microglia signature from the current GTEx and ABA tissue data. The derived signature included all known markers of microglia, including *TMEM119, P2RY12, and CD68* (Bennett et al., 2016; Perego, Fumagalli, & De Simoni, 2011; Wes, Holtman, Boddeke, Möller, & Eggen, 2016) and many other genes known to be associated with microglial/macrophage biology. This includes representatives of the TLR, complement, and MHC class 2 antigen-presenting immune pathways.

Validation of the signature, included an examination of HPA immunostaining of proteins encoded by the signature genes, demonstrating that they were significantly expressed in myeloid cell types relative to other neuronal and immune cells, and comparison with published microglial signatures. This final step revealed approximately half of the core genes as conserved between species, reaffirming the widely accepted idea that many constitutively expressed microglial genes are conserved between mouse and human. Although their response to processes like aging may diverge (Bennett et al., 2016; Galatro et al., 2017). To examine the specificity of the core signature, a comparison was made with the Galatro *et al.* (Galatro et al., 2017) signature, derived by comparing the fold expression of microglia isolated from the parietal cortex of post-mortems, relative to whole tissue. The signature provides an insight into human microglial functionality under homeostasis and has a high degree of overlap with current signatures, although being significantly larger. Using the GTEx brain atlas data the coexpression and regional expression of both signatures were investigated. Many of the Galatro *el al.* signature genes showed poor coexpression while displaying a range of region-specific expression patterns, deviating from those of canonical marker genes like CSF1R, AIF1 and CX3CR1. Whilst these genes are likely to be expressed in microglia (as originally identified), these results underscore the regional heterogeneity of microglia, suggesting that certain genes specific to cortical microglia may not be solely expressed by microglia in other regions. Indeed, certain of the Galatro signature genes, also common to other studies, expressed highly in the cerebellum presenting a multi-cell type expression in this region based on IHC data from the HPA, making them poor markers of microglia. Additionally, these genes did not agree with cerebellum-specific genes identified in mouse (Grabert et al., 2016). In contrast, our core signature exhibited a well-defined and condensed coexpression pattern corresponding to the known regional CNS variation in microglial abundance. Therefore, while isolated cells have provided fundamental insight into microglial identity, coexpression analysis of the employed datasets aids in defining the microglial specific profile in the CNS.

Evidence for the central role microglia play in the pathogenesis of neurodegenerative disease continues to grow, however, the cellular and molecular changes that occur in human brain pathologies are poorly understood. Furthermore, using the core signature we conducted various analysis, to discern the influence of cell number and activation state in both healthy aging and Alzheimer’s, across a number of brain regions using the dataset generated by Berchtold *et al.* (Berchtold et al., 2013). Given that the majority of microglial genes maintain their expression with age (Galatro et al., 2017; Jyothi et al., 2015; Poliani et al., 2015), we made the assumption that the average expression of the signature genes, can be used as a proxy for microglia number through aging. Supporting this, the increased average expression of signature genes with age was substantiated by studies directly measuring cell numbers with age (Damani et al., 2011; Peters, Josephson, & Vincent, 1991; Tremblay, Zettel, Ison, Allen, & Majewska, 2012). The largest changes were observed in the hippocampus, a region particularly vulnerable to aging and where greater microglial activation and neuronal loss have been observed with age relative to other cortical regions (Bartsch & Wulff, 2015; Galatro et al., 2017; Kumar et al., 2012; Raz et al., 2005). Therefore, these analyses support the idea that microglial numbers change in a region-dependent manner and that these changes correlate with age-associated regional atrophy and inflammation. When comparing Alzheimer’s to age-matched controls, a similar trend towards increased expression levels of signature genes was also observed. However, a significant increase was only observed in the superior frontal gyrus, a region known to be highly susceptible to the effects of both aging and Alzheimer’s, based on neuronal connectivity studies (Bakkour et al., 2013; Stam, 2014). Interestingly, the entorhinal cortex and hippocampus, whose atrophy characterize Alzheimer’s pathology, showed the greatest differences between Alzheimer’s and controls, although lacking statistical significance, likely due to the relatively small number of samples and large variability between them (Khan et al., 2014; Velayudhan et al., 2013). In contrast, the postcentral gyrus, a region shown to maintain its grey matter content and functional connectivity with other regions in late-onset Alzheimer’s, showed little change in Alzheimer’s versus controls (Adriaanse et al., 2014; Thompson et al., 2003). Strikingly, these findings are consistent with regional Alzheimer’s progression based on tau burden, neuroinflammation and neuronal loss, which are prominent in the entorhinal cortex and hippocampus (Cope et al., 2017; Freer et al., 2016; Kreisl et al., 2016). Overall, these data demonstrate the utility of the signature in assessing quantitative differences in microglial numbers from tissue-level expression datasets.

To gain insight into molecular pathways specifically affected in Alzheimer’s, qualitative changes in the profile of microglia were examined. Coexpression analysis identified a set of 165 MAGs correlating with the core gene signature in samples isolated from Alzheimer’s patients. The MAG list was enriched in various pathways associated with innate immune signaling, consistent with the inflammatory environment within Alzheimer’s brain tissue and the reactivity of microglia within this environment, namely *TSPO* (Kumar et al., 2012). It was particularly interesting to note that genes involved in lipid regulation and wound healing, associated with Alzheimer’s, were over-represented in the MAGs set (Cervantes et al., 2011; Lorenzl et al., 2003; Petit-Turcotte et al., 2001; Y.-H. Shih et al., 2014). Members of the *APOC* gene family and *ECHDC3* are known to regulate levels of certain lipids, linked with Alzheimer’s progression (Adunsky et al., 2002; Desikan et al., 2015; Lane & Farlow, 2005). Additionally, these factors are part of the wound healing cascade, including proteins such as *TIMP1* and *PROS1*, which are key in regulating tissue integrity and plasticity, altogether pointing towards the vulnerable blood-brain barrier in Alzheimer’s (Bennett et al., 2016; Duits et al., 2015). These results provide some insight and support for the complexity of microglia involvement in Alzheimer’s through not only inflammatory mechanisms but also through upregulation of metabolic and tissue homeostasis/repair functions (Vincenti et al., 2016). Investigating quantitative alterations of microglial differentiation in Alzheimer’s, we focussed on genes differentially expressed in Alzheimer’s compared to age-matched controls. Although for all regions, the majority of genes presented an upward trend of expression, most lacked significance, excluding those of the superior frontal gyrus which we further investigated. Genes relating to TYROBP signaling, which is implicated in Alzheimer’s and together regulates phagocytosis, cell proliferation, activation and survival were significantly upregulated (Keren-Shaul et al.; Landreth & Reed-Geaghan, 2009; Ma et al., 2015). Substantiating these findings TYROBP knockout mice models have proven to suppress inflammation in neurodegenerative models including Alzheimer’s, thereby minimizing neuronal dystrophy, implicating a failure in the resolution of inflammation in Alzheimer’s (Bakker et al., 2000; Haure-Mirande et al., 2017). Interestingly mutations and expression of downstream members are also linked with Alzheimer’s including CD33, TREM2 and CR3 (Hamerman, Tchao, Lowell, & Lanier, 2005; Takahashi, Rochford, & Neumann, 2005).

In summary, we have employed a coexpression analysis approach to derive a core human microglial signature under non-neuropathologic conditions that is robust to potential artifact generated by technical and biological variation (e.g. donors and CNS regions) that can influence other approaches in signature derivation. Furthermore, we present the utility of this signature, demonstrating its sensitivity to detect region-specific changes in microglial alterations in aging and Alzheimer’s disease, while appreciating the influence of cell numbers and activation in tissue transcriptomics data. We found that these responses were aligned with the known neuropathological trajectory of Alzheimer’s. We propose the conserved signature described here as a specific and robust resource of gene markers that reflect the core functional profile of these cells and aid future studies of microglial biology in the human CNS, including bulk and single cell transcriptomics.

## Acknowledgments

T.C.F. and B.W.M. are funded by an Institute Strategic Programme Grant funding from the Biotechnology and Biological Sciences Research Council [BB/J004227/1]. B.W.M. receives funding from the UK Dementia Research Institute and Medical Research Council [MR/L003384/1]. B.S. is supported by Experimental Medicine Challenge Grant funding from the Medical Research Council [MR/M003833/1].

---

Table S1. Metadata of datasets used in the analysis.

Table S2. Dataset derived microglial clusters

Table S3. Core human microglial signature with annotation

Table S4. ToppGene enrichment analysis of core microglial genes

Table S5. Clusters from figure 4

Table S6. Fold change in core and MAG for aging and Alzheimer’s across regions

Table S7. ToppGene enrichment analysis of MAG.

Table S8. ToppGene enrichment analysis of genes differentially expressed in Alzheimer’s and not in aging.

**Figure S1.**
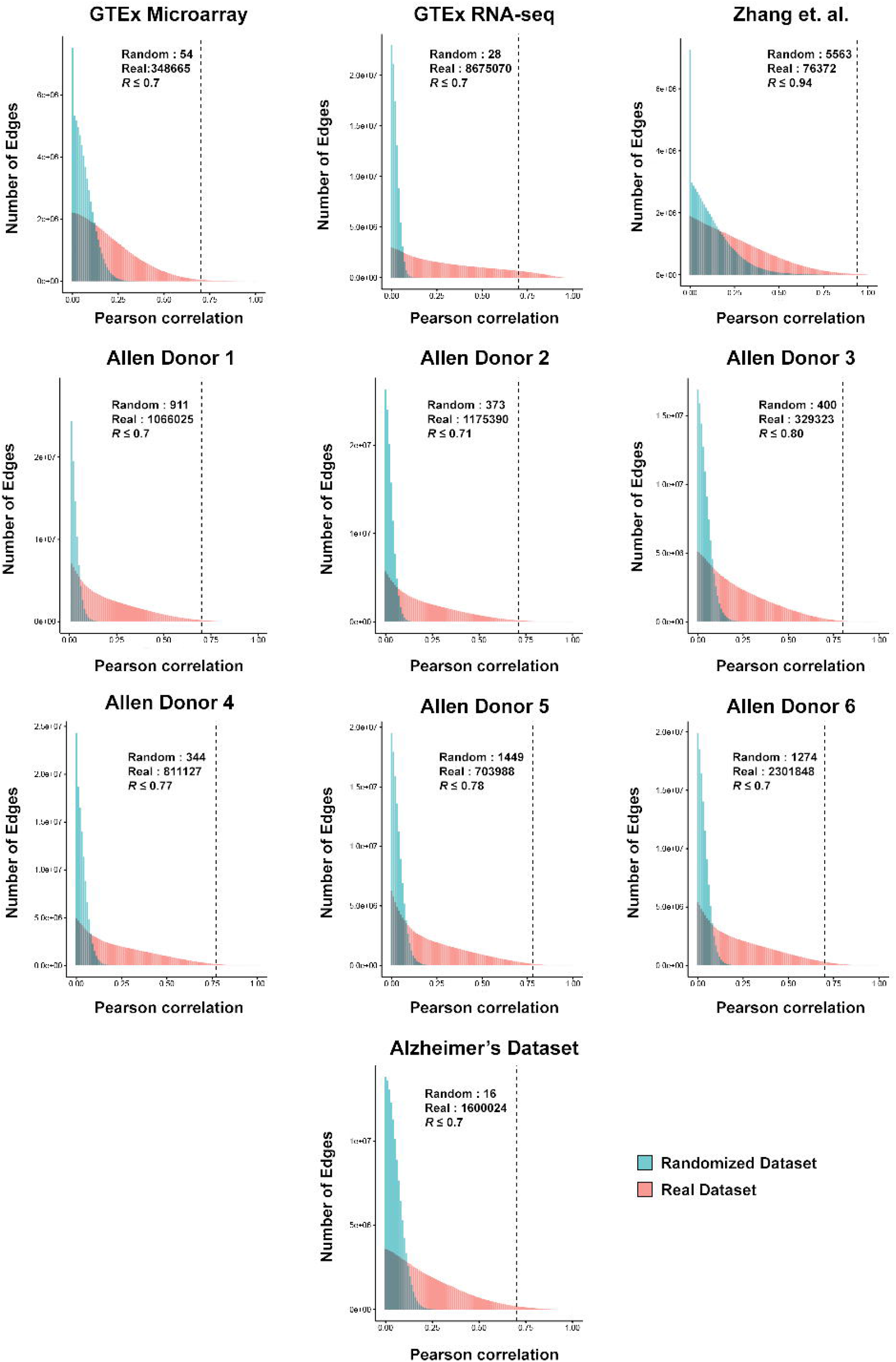
Distribution of Pearson correlations for each dataset analyzed. Plots show the distribution of positive Pearson correlations (edges) between genes (nodes) for Pearson correlations between 0 and 1 observed for each dataset (red), relative to the distribution of correlations for the equivalent randomized dataset (blue). The dotted line shows correlation threshold used for analyses and figures quoted list the number of edges one would expect by chance (random) versus those observed (real).

**Figure S2.**
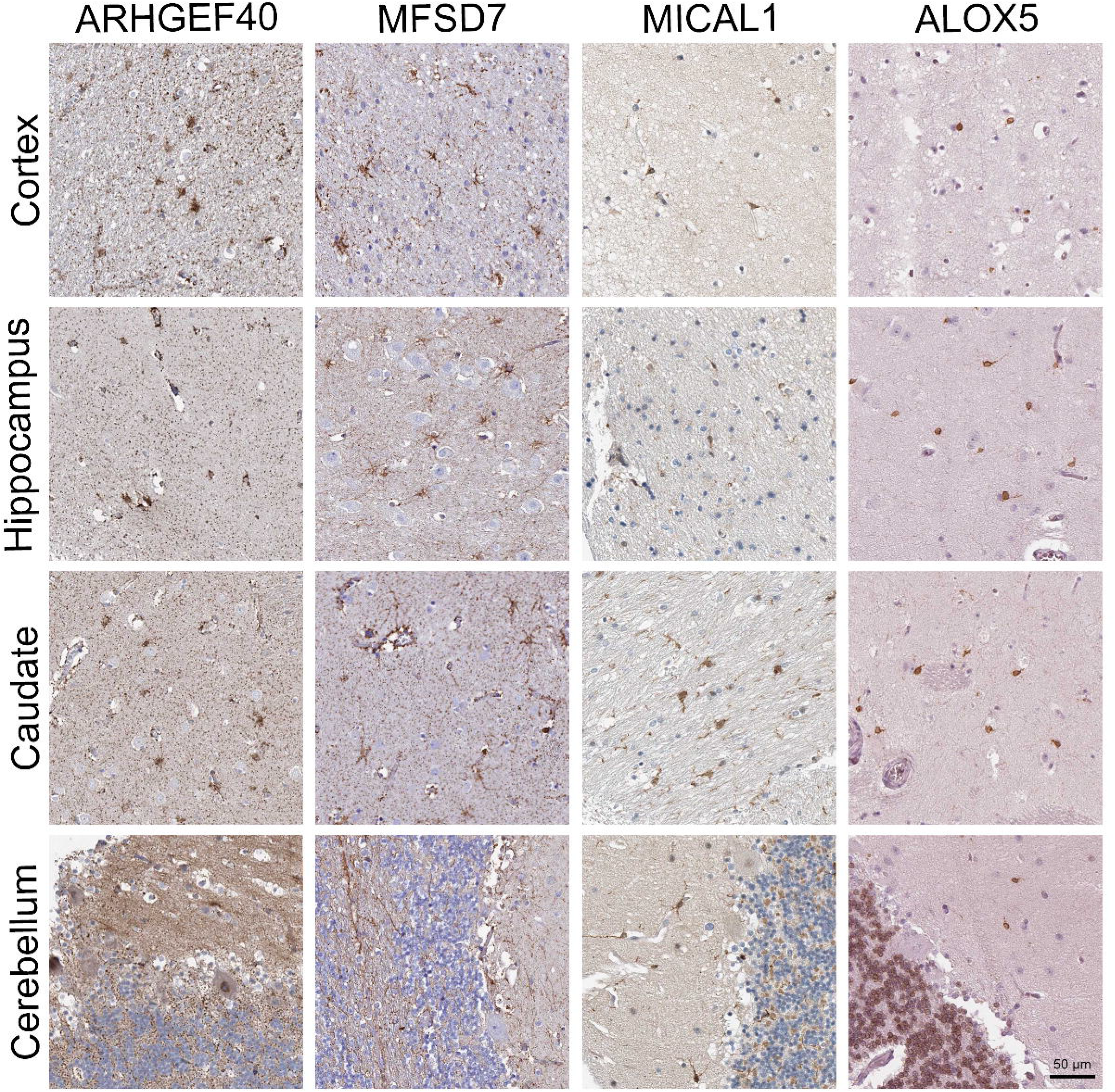
IHC of Galatro *et al.* signature genes coexpressed in the cerebellum. IHC staining for a number of proteins from the Galatro et al. gene signature found to be significantly expressed in the cerebellum relative to other regions. IHCs were taken from the HPA resource (Nilsson et al., 2005), and show that in the cerebellum their expression is not restricted to microglia.

